# Tumor γδ T-cell abundance is associated with favorable cancer treatment outcomes

**DOI:** 10.64898/2026.08.27.747587

**Authors:** Xingzhi Niu, Deepali Kundnani, Mikaela Dicome, Luis Tafoya, Li Song, Murad Mamedov, X. Shirley Liu, Avinash D. Sahu

## Abstract

**Purpose:** Clinical response to immune checkpoint blockade (ICB) remains variable. We performed a reference-wide screen to identify immune-cell populations in the tumor microenvironment (TME) associated with benefit across treatment modalities and tumor types.

**Experimental Design:** We analyzed pretreatment bulk tumor RNA-seq to identify RNA-abundance patterns associated with favorable outcomes in (1) published ICB cohorts with response labels and (2) pretreatment TCGA cohorts with survival outcomes. To identify associated cell types, we compared favorable expression patterns to previously determined patterns for 157 cell types defined by the Human Primary Cell Atlas (HPCA), spanning immune, stromal, vascular, and epithelial lineages. Cell types were associated with favorable outcomes using Cox models (TCGA survival) and mixed-effects models of responder status that accounted for cohort, therapy class, tumor type (ICB cohorts). ICB analyses were restricted to pretreatment samples. Random-effects meta-analysis was used for the PRECOG cross-platform replication. After γδ T-cells emerged as the top-ranked population associated with ICB response, we adjusted γδ associations for CD8^+^ T-cell abundance, and evaluated γδ signals with TRUST4-based T-cell receptor gamma (TRG) and delta (TRD) gene reconstruction and single-cell RNA-seq. To test whether ICB-favorable cellular landscapes were shared with non-immunotherapy treatment response, we used TCGA RECIST-evaluable treatment records to derive immune-cell profiles associated with treatment response for chemotherapy, radiation, targeted therapy, and hormone therapy.

**Results:** The γδ T-cell program was the top-ranked cell-type program associated with ICB response and was also among the programs most strongly associated with favorable TCGA survival. Cell types associated with ICB response showed strong descriptive concordance with the chemotherapy-response profile derived from recorded TCGA outcomes (Pearson r of ranks = 0.92) and moderate concordance with the analogous radiation-associated profile (r = 0.58). Analyses of targeted and hormone therapies were limited by small sample sizes and showed weaker concordance. In pretreatment tumor biopsies, inferred γδ T-cell abundance was associated with ICB response and with survival, and these associations were not explained simply by CD8^+^ T-cell abundance. In TCGA, TRUST4-derived γδ T-cell receptor (TCR) abundance estimates correlated with signature-based γδ scores and stratified survival. In independent single-cell RNA-seq datasets, higher γδ TCR RNA abundance was also associated with a higher rate of ICB response.

**Conclusion:** A reference-wide cell-type screen identified γδ T-cells as the leading population associated with ICB response, and pretreatment γδ T-cell abundance was also associated with favorable outcomes in immunotherapy and non-immunotherapy settings. The evidence is associative and varies in strength by tumor context but supports prospective evaluation of γδ T-cell abundance as a component of pretreatment immune profiling alongside CD8, B-cell/TLS, stromal, and epithelial-state features.

**Translational Relevance:** Across cancers, clinical response to immune checkpoint blockade (ICB) is variable, with inconsistent performance from established biomarkers such as PD-L1 and tumor mutation burden. We asked which immune-cell transcriptional programs were associated with favorable outcomes across immunotherapy and non-immunotherapy treatment settings. Using pretreatment bulk tumor RNA-seq from published ICB cohorts and TCGA, we evaluated the inferred abundance of 157 subtypes in the tumor microenvironment (TME) with a common gene-level and sample-level framework. Pretreatment TMEs enriched in lymphoid cells or γδ T-cells were associated with ICB response and with favorable survival. The immune-cell landscape associated with favorable ICB outcomes was strongly concordant with chemotherapy response and moderately concordant with radiation response. The association between γδ abundance and favorable outcomes remained after accounting for CD8^+^ T-cell abundance. These observational findings support γδ T-cell abundance as a candidate for pretreatment immune profiling that warrants prospective evaluation.

## 1 Introduction

Immune checkpoint blockade (ICB) releases inhibitory brakes on T-cells, most notably via the PD-1/PD-L1 and CTLA-4 pathways (1). The canonical model of ICB efficacy relies on cytotoxic CD8^+^ T-cells recognizing tumor neoantigens presented on MHC Class I molecules. While ICB has transformed oncology, this model fails to explain the full spectrum of clinical observations. Many tumors evade immune surveillance by eliminating expression of MHC Class I (2), yet some of these “invisible” tumors still respond remarkably well to ICB therapy. A striking example is classical Hodgkin lymphoma (cHL), where response rates to PD-1 blockade approach 70% despite malignant cells frequently lacking MHC Class I expression. This paradox strongly suggests that the antitumor immune response extends beyond the classical MHC-restricted CD8^+^ T-cell axis (3, 4).

Across cancers, large-scale analyses have shown that immune lineages other than CD8^+^ T-cells shape clinical outcomes. Pan-cancer and multi-cohort studies have linked diverse leukocyte subsets, particularly B cells and tertiary lymphoid structures (TLS), to better prognosis under standard therapies and to ICB response (5, 6). Collectively, these studies indicate that cytotoxic CD8^+^ T-cells (7, 8) work in conjunction with B cells, TLS, and γδ T-cells to promote favorable outcomes in selected tumor types (5, 6, 9–11). Unlike αβ T-cells, γδ T-cells can recognize intact proteins, lipid antigens, phosphoantigens, and stress-induced molecules directly, without dependence on classical MHC antigen presentation (9–11).

Most existing γδ T-cell studies are limited to single tumor types or small cohorts, and ICB and non-ICB settings are often analyzed separately. It remains unclear whether immune-cell programs associated with ICB response are the same as those associated with favorable outcomes under other treatments. The clinical implications of γδ infiltration seem to be context dependent. Some studies report favorable associations in dMMR tumors, Merkel cell carcinoma, cervical cancer, and breast cancer (1, 12–15), whereas others report weaker or adverse associations, including in some head and neck squamous cell carcinoma cohorts (16). A further unresolved question is whether γδ T-cell abundance is simply a feature of a generally inflamed, CD8-rich microenvironment, or whether it has significance on its own.

To address these questions, we analyzed the inferred abundance of 157 pre-defined cell types and their subtypes across ICB and TCGA bulk RNA-seq datasets. First, we identified immune-cell populations associated with ICB response and with TCGA survival, testing whether these associations converged on a shared favorable microenvironment. Second, we asked whether the ICB-favorable cell-type landscape was also observed in TCGA responses to chemotherapy, radiation, targeted therapy, and hormone therapy. Third, we tested whether γδ T-cell abundance, a top candidate, remained associated with outcome after adjustment for CD8^+^ T-cell abundance and whether orthogonal approaches such as T-cell receptor gamma (TRG) and delta (TRD) gene reconstruction and single-cell RNA-seq supported the conclusions drawn from bulk-RNA signals.

## 2 Patients and Methods

### 2.1 Patients and cohorts

#### ICB cohorts

We used the RNA-seq and response calls from the following published cohorts: Gide *et al.* 2019 (melanoma; anti-PD-1 monotherapy and anti-PD-1+anti-CTLA-4) (17); Hugo *et al.* 2016 (melanoma; anti-PD-1) (18); Liu *et al.* 2019 (melanoma; PD-1 blockade) (19); Mariathasan *et al.* 2018 (urothelial carcinoma; anti-PD-L1) (20); Miao *et al.* 2018 (microsatellite-stable solid tumors; ICB) (21); Nathanson *et al.* 2017 (melanoma; anti-CTLA-4) (22); Riaz *et al.* 2017 (melanoma; nivolumab) (23); Van Allen *et al.* 2015 (melanoma; anti-CTLA-4) (15); Zhao *et al.* 2019 (glioblastoma; anti-PD-1) (24); Braun *et al.* 2020 (renal cell carcinoma, anti-PD-1) (25); Campbell *et al.* 2023 (melanoma; anti-PD-1+anti-CTLA-4) (26); McDermott *et al.* 2018 (renal cell carcinoma; anti-VEGF, anti-PD-L1, and anti-VEGF+anti-PD-L1) (27); Kim *et al.* 2018 (gastric carcinoma; anti-PD-1) (28); Uppaluri *et al.* 2020 (head and neck squamous cell carcinoma; anti-PD-1) (29); Lee *et al.* 2021 (non-small cell lung cancer; anti-PD-1) (30). All post-treatment biopsies were excluded from the RNA-seq analyses. All analyses were performed on pretreatment tumor RNA-seq only.

#### TCGA cancers (non-immunotherapy)

We analyzed TCGA RNA-seq, cancer type, survival outcomes, and, where specified, GDC treatment records with RECIST measure_of_response annotations (31). TCGA survival analyses used cancer-type-stratified or cancer-type-adjusted models. Treatment-modality response analyses used only RECIST-evaluable treatment lines and excluded patients with any documented immunotherapy exposure.

### 2.2 Data processing

Gene-level RNA-seq was harmonized within each cohort (ICB) or cancer type (TCGA). Counts were converted to TPM and restricted to protein-coding genes. Low-expression genes were excluded. For any continuous analysis, values were z-standardized within cohort or within cancer type to remove scale differences and facilitate cross-study modeling. The z score was calculated by subtracting the within-set mean from each TPM value and dividing by the within-set standard deviation:

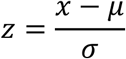

### 2.3 Immune reference and sample-level abundance

The HPCA, a curated reference compendium that defines 157 cell types and subtypes from purified bulk and single-cell resources (32), was used for (1) projection of gene-level outcome effects onto reference cell types, and (2) indirect estimation of cell-type abundance for each tumor sample by correlating each tumor transcriptome with reference profiles. Before modeling association with treatment response or survival, these tumor-sample scores were z-standardized within each ICB cohort or TCGA cancer type.

### 2.4 Analytic modules

#### ICB differential expression

For each gene j, we fit a linear mixed-effects model across pre-treatment ICB samples:

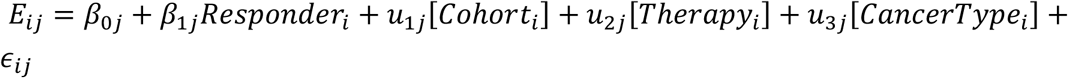

Here, E*_ij_* is the normalized expression of gene *j* in sample *i*, and *Responder_i_* indicates clinical response to ICB (1 = responder). The coefficient β*_1j_* estimates the responder-associated expression effect for gene *j* after accounting for cohort, therapy class, and cancer type. Cohort, therapy, and cancer type were modeled as random intercepts.

#### Non-ICB Cox mixed-effect model

For each gene, we fit a Cox proportional-hazards model with a cancer-type random intercept (33) to relate expression to overall survival (OS) or progression-free interval (PFI):

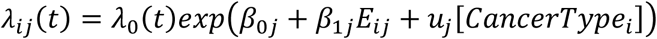

Here, λ_0_(*t*) is the baseline hazard and β_1*j*_ estimates the association between expression of gene *j* and survival. The cancer-type random intercept accounts for baseline outcome differences across TCGA projects.

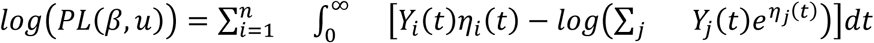

*Y_i_*(*t*) denotes whether subject *i* remains under observation at time *t*, and η*_i_*(*t*) is the linear predictor. Fixed and random effects are represented by design matrices X and Z, respectively.

#### Treatment-modality response, differential expression (TCGA)

To compare the ICB-response cell landscape with response landscapes for specific non-ICB treatments, we built per-gene responder-vs-non-responder differential-expression analyses for chemotherapy, radiation, targeted therapy, and hormone therapy using TCGA cases with documented treatment records and RECIST “measure_of_response” annotations. Treatment lines were classified by keyword matching of “therapeutic_agents” and “treatment_type” fields. Response was binarized within each modality as completed response (CR) or partial response (PR) versus progressive disease (PD) or stable disease (SD). Patients with any documented immunotherapy exposure were excluded from the four non-ICB modality analyses. We used TCGA primary-tumor RNA-seq TPM, log2(x+1)-transformed and z-scored per gene within cancer type. For each gene, we fit a weighted linear model, expression ∼ response + cancer + project, with weights of 1 / number of treatment lines per patient. Projects with fewer than two responders or two non-responders were excluded. The response t-statistic for each gene was projected onto HPCA (Human Primary Cell Atlas) cell types as described in section 2.3, and concordance with the ICB-response profile was computed across the fine-grained 157 HPCA cell types (“label.fine” annotations).

#### Cell-type-level differential abundance

We z-scored each gene across the 157 reference populations and projected gene-level outcome effects onto each reference profile by Spearman correlation. Positive projection scores indicate that genes characteristic of a reference cell type align with expression patterns associated with ICB response or favorable TCGA prognosis; negative scores indicate the opposite. When a biological cell type had multiple reference profiles, each profile yielded a separate projection score.

#### Sample-level immune abundance estimation

We estimated per-sample immune features, including γδ T and B-cell/TLS scores, by correlating each pretreatment tumor transcriptome with the 157-cell-type reference using a SingleR-inspired (32) bulk RNA-seq adaptation. Scores were z-standardized within ICB cohort or TCGA cancer type. We benchmarked these scores in three ways: (i) pseudo-bulk simulations with known cell fractions: recovery was quantified as the Spearman correlation between each sample’s score and its true fraction, computed against an independently seeded reference (different from the data generator) so that recovery is non-circular; γδ and CD8 fractions were recovered at ρ = 0.73 and 0.81 (mean over 8 replicates; Fig. S2a and Fig. S2b), with performance declining gradually under added noise (Fig. S2c) and varied mixture composition (Fig. S2d); (ii) concordance with established deconvolution tools across ICB cohorts; and (iii) comparison with IMvigor210 immunohistochemistry-defined tumor immune phenotypes, namely “immune-inflamed”, “immune-excluded”, and “immune-desert” tumors.

#### CD8-adjusted outcome models

To evaluate whether γδ abundance carried information beyond CD8-associated inflammation, we fit γδ outcome models with CD8 covariates. In TCGA, cancer-type-specific Cox models included z-standardized γδ and CD8 scores. In the integrated ICB analysis, γδ response was modeled with a binomial mixed-effects logistic model (random intercept per cohort) and overall survival with a cohort-stratified Cox model; each model was refitted after adding one of the eight CD8 estimators — our correlation-enrichment CD8 score, CD8A/CD8B mean expression, CIBERSORT, EPIC, TIMER, xCell, MCP-counter, and quanTIseq — as a covariate, with γδ and CD8 z-scored within cohort (Fig. S3). In the single-cohort IMvigor210 analysis (20), γδ response (logistic) and OS (Cox) models were each additionally adjusted, one at a time, for the eight CD8 estimators and for the IHC immune-phenotype category (inflamed/excluded/desert). As a specificity control, each CD8 estimator was also fit as the predictor in place of γδ, alone and with the IHC phenotype (Fig. S4).

#### TCR reconstruction and single-cell RNA-seq support

We reconstructed TRG/TRD CDR3s using TRUST4 (34), computed a TRUST4 γδ score, and analyzed TRUST4 outputs within each TCGA cancer type. We also evaluated expression of γδ-associated TCR genes (*TRGV*, *TRDV*, *TRDC*) in published immunotherapy single-cell RNA-seq datasets from patients with categorized responses. These analyses provide orthogonal support for the bulk γδ signal.

#### Pipeline calibration by simulation

To assess detection sensitivity (power) and type-I error of the pipeline, we generated pseudo-bulk RNA-seq cohorts by Dirichlet-mixing the reference cell-type profiles (10 cohorts × 60 samples per replicate) under four scenarios: we simulated response as depending on CD8 abundance (S1), γδ abundance (S2), both (S3), or no cell type (S4, the global null), at prespecified effect sizes (log-odds per SD). Samples were scored with cell-type-specific marker signatures against an independently seeded reference (so recovery is non-circular) and z-scored within cohort. Because some signatures are correlated — γδ and CD8 share marker genes — all cell types were estimated jointly in a single multivariable logistic model (sample response on all cell-type scores plus a cohort term); each cell type’s significance, taken from its own coefficient, is therefore conditioned on the others rather than confounded by them. Power was the proportion of replicates (n = 12 per effect size, 0.25–1.5 SD; the 2.0-SD point was omitted owing to logistic saturation) in which the planted cell type reached p < 0.05; type-I error was the proportion of 50 null replicates in which a cell type reached p < 0.05. A marginal, one-cell-at-a-time version of the same test inflated the γδ false-positive rate to ∼0.10, whereas the joint model held it at the nominal level (0.04 at α = 0.05), motivating the joint specification reported in Fig. 1E and Fig. S1. This pipeline calibration is distinct from the per-sample abundance-score validation (Fig. S2a--d).

**Fig. 1:**
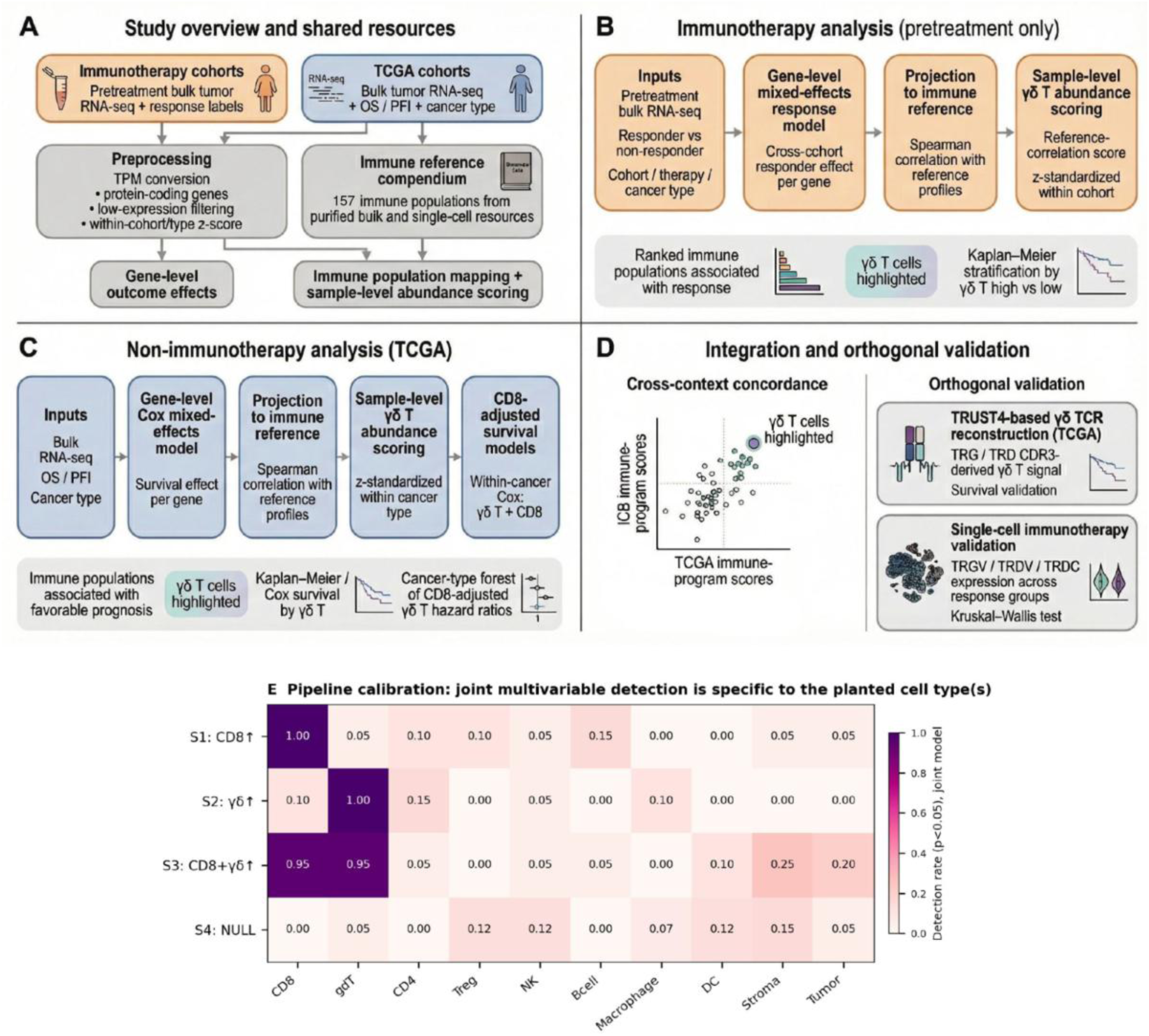
Study overview and analytic framework for identifying γδ T-cell-associated immune programs across immunotherapy and non-immunotherapy contexts. (A) Pretreatment ICB RNA-seq, TCGA RNA-seq, and a 157-cell-type reference were harmonized for gene-level and sample-level analyses. (B) In pretreatment ICB cohorts, mixed-effects responder models were projected onto the immune reference to rank cell programs associated with response. (C) In TCGA, Cox survival models were projected onto the same reference, followed by sample-level γδ T scoring and CD8-adjusted survival analysis within cancer type. (D) Cross-context analyses compared ICB and TCGA cell-type axes and tested the γδ T-cell signal with TRUST4-based TCR reconstruction and single-cell immunotherapy data. (E) Planted cell types are shown on the y-axis: CD8, γδ, joint CD8 + γδ, and null. Other immune cell type signatures are shown on the x-axis. Under a joint multivariable model of cell-type-specific signatures, effects are detected specifically for the planted cell type while type-I error is held at the nominal level (see also Fig. S1).

#### TCGA-PRECOG cross-platform concordance

To test whether prognostic cell-state effects transferred across expression platforms, we applied the same HPCA correlation-enrichment scorer to TCGA RNA-seq and 108 evaluable PRECOG (5) microarray cohorts. Cox OS models used within-cohort z-scored cell-type scores; PRECOG cohorts were pooled within cancer type by DerSimonian-Laird random-effects meta-analysis (35). We compared matched PRECOG-TCGA cancer pairs using the vector of cell-level log-HRs across the 157 reference profiles and, separately, the prespecified γδ score. As prespecified before examining the results, cohorts with fewer than 30 patients were excluded from the analysis. Per-cohort Cox models were retained only when they converged to a finite log-hazard ratio.

### 2.5 Multiple testing, assumptions, and sensitivity analyses

Unless otherwise noted, all tests were two-sided, with Benjamini-Hochberg FDR control within analysis families. Proportional-hazards assumptions were checked where Cox models were used, and sensitivity analyses included alternative γδ estimation, CD8 adjustment, simulation-based pipeline calibration, and TCGA-PRECOG cross-platform concordance.

### 2.6 Software and reproducibility

Analyses were performed in R (version 4.3 or later) and Python. Key packages and tools included lme4/lmerTest for mixed-effects differential expression, survival and coxme for survival models, metafor for random-effects meta-analysis, TRUST4 for repertoire reconstruction, and ggplot2 and ggrepel, or matplotlib for visualization. Figure-source data tables and code will be deposited with the public repository at publication.

### 2.7 Ethics, data, and code availability

All data were obtained from public, de-identified sources (see section 2.1). No new patient enrollment occurred. Data use in this study complied with original study terms. Code and derived data will be made available in a public repository upon publication, in accordance with CCR/AACR policies.

## 3 Results

### 3.1 A unified pan-cancer framework links tumor immune-cell programs to clinical outcome

To identify tumor-infiltrating immune populations associated with favorable clinical outcomes, we used a common pipeline to analyze (1) pretreatment ICB cohorts with response labels and (2) TCGA tumors with survival outcomes, including overall survival (OS) and progression-free interval (PFI). Across the ICB cohorts, gene-level outcome models estimated responder-associated expression, whereas, for TCGA tumors, the models estimated survival-associated expression. We then projected these gene-level effects onto the curated 157-cell-type HPCA reference, which includes immune, stromal, vascular, and epithelial lineages, while accounting for cohort, therapy class, and cancer type (Fig. 1A-C).

Because the same immune reference was used in both settings, each cell-type profile could be compared across ICB response and TCGA survival. We used these ranked cell-type profiles to identify shared favorable programs. We then prioritized the top candidate, γδ T-cell abundance, for patient-level follow-up with bulk RNA-seq scoring, CD8 adjustment, TRUST4-based TRG/TRD reconstruction, and single-cell immunotherapy datasets (Fig. 1D).

We first used simulation to test two properties of the pipeline: whether it recovered true cell-type effects (sensitivity) and controlled spurious effects (type-I error) (Methods). Because γδ and CD8 signatures share marker genes, all cell types were estimated together in a single joint multivariable model, so each cell’s effect is conditioned on the others. When a γδ effect was introduced into the simulated cohorts, the pipeline recovered it at ∼100% power across realistic effect sizes, while a non-planted CD8 signature stayed near the chance level (Fig. S1a); the joint model recovered each planted population specifically, leaving the others at the nominal rate (Fig. 1E). Under a global null in which no cell type was associated with outcome, the γδ false-positive rate stayed at the nominal level (0.04 at α = 0.05), with no γδ-specific inflation (Fig. S1b).

Separately, we confirmed that the per-sample scores reflected cell abundance. Against simulated mixtures with known cell fractions, the pipeline recovered the true γδ and CD8 T-cell fractions (Spearman ρ = 0.73 and 0.81; Fig. S2a and Fig. S2b), and recovery remained positive as noise (Fig. S2c) and mixture composition varied (Fig. S2d).

### 3.2 ICB responders have an interferon-rich, cytotoxic, lymphoid-rich pretreatment state

We asked which pretreatment transcriptional programs distinguished ICB responders from non-responders. Across the pretreatment ICB RNA-seq cohorts, responders showed higher expression of genes linked to interferon signaling and antigen presentation (*IFNG*, *IRF1*, *TAP1*), cytotoxic lymphocyte effectors and recruitment chemokines (*NKG7*, *GZMA*, *CCL5*, *XCL2*, CXCL9), and T-cell activation or differentiation (*ZNF683*, *LAG3*) (Fig. 2A-B). This pattern is consistent with an “immune-inflamed” pretreatment state being favorable to ICB response.

**Fig. 2:**
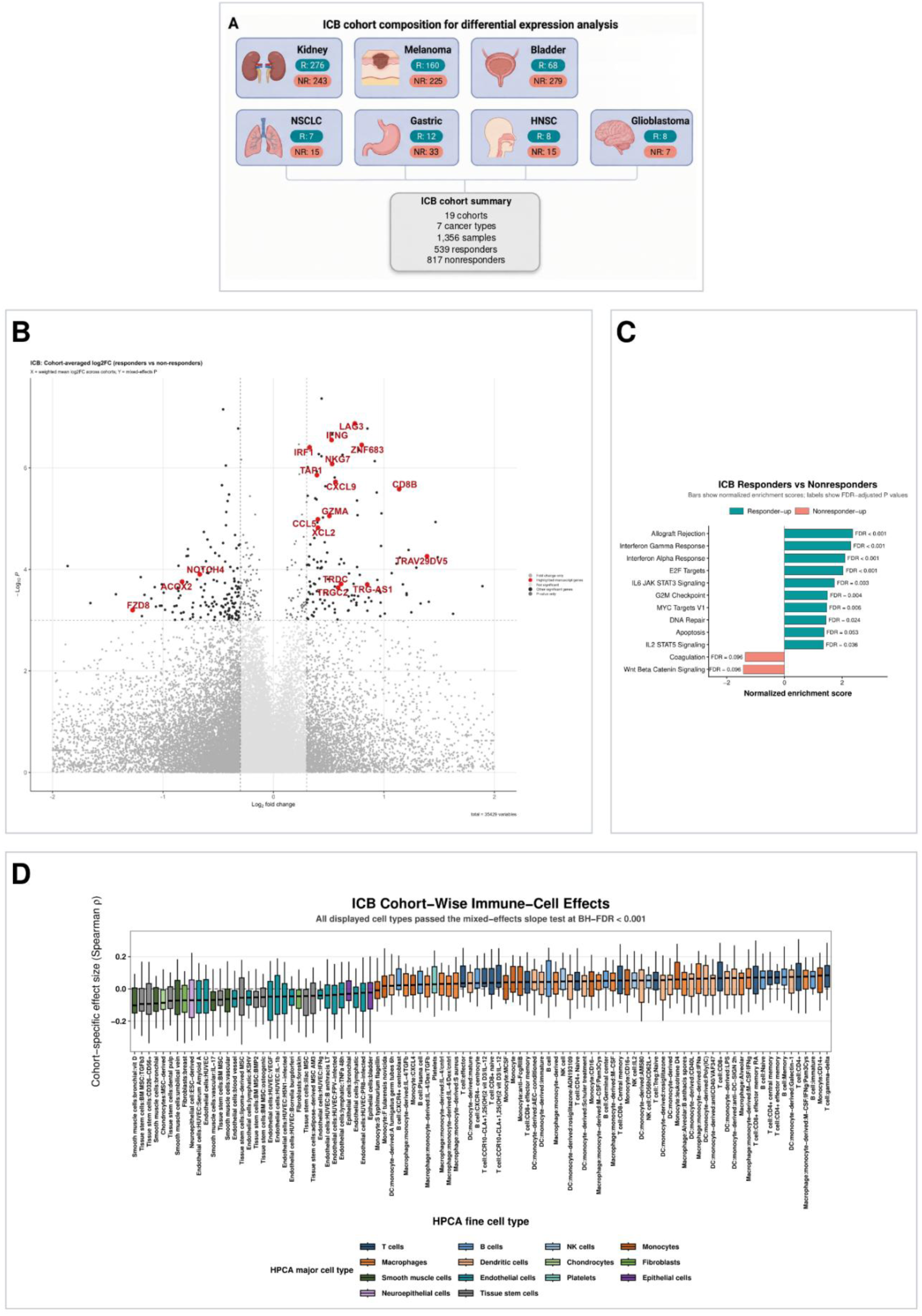
Pretreatment ICB cohort composition and cross-cohort differential-expression analysis identify responder-associated immune programs. (A) Composition of the pretreatment ICB cohorts used for responder-versus-non-responder analysis. (B) Mixed-effects volcano plot showing genes differentially expressed between responders and non-responders. (C) Hallmark pathway enrichment showing immune and inflammatory pathways enriched in responders and WNT/β-catenin plus coagulation enriched in non-responders. (D) Cohort-wise projection of responder-versus-non-responder gene-expression effects onto HPCA reference profiles. For each cell population, the central black line denotes the median cohort-specific Spearman projection score (ρ), the box spans the interquartile range, with positive scores indicating alignment with response-associated expression.

In responders, pathway enrichment analysis identified IFN-α and IFN-γ response, allograft rejection, IL2/STAT5, IL6/JAK/STAT3, E2F, G2M, MYC, and DNA-repair programs. By contrast, in non-responders, WNT/β-catenin and coagulation programs were enriched (Fig. 2C). This analysis connects response with antigen presentation, effector recruitment, and proliferative immune activation, while non-response aligns more closely with immune-exclusion, wound healing, and coagulation-associated biology.

We then projected the responder differential-expression signature onto immune-cell reference profiles. Lymphoid populations showed a positive shift in the projection score (toward responder enrichment), consistent with enrichment in responders, whereas mesenchymal–stromal programs (MSC/chondrocyte, fibroblast, and endothelial states) have negative projection scores, consistent with relative depletion in responders. Within the lymphoid compartment, the γδ T-cell program showed strongest association with response (Fig. 2D). This analysis defines a pretreatment ICB responder state in which interferon signaling, antigen presentation, cytotoxic effectors, and lymphoid enrichment co-occur. γδ T-cells emerged from within that broader state rather than as an isolated marker.

### 3.3 A lymphoid-rich pre-treatment TME is associated with better prognosis among TCGA samples

The preceding ICB analysis ranked all 157 cell types according to their association with treatment response and provides an opportunity for comparison of associations between TME cell-type composition and outcomes across treatments. We therefore applied the same framework to TCGA to ask whether related lymphoid biology was associated with survival in other cancer cohorts. Among TCGA tumors with evaluable survival data, the pan-cancer gene-level Overall Survival (OS) landscape separated into favorable and unfavorable tails (Fig. 3A-B). The unfavorable tail was enriched for proliferation, stromal remodeling, epithelial-mesenchymal transition, invasion, and metabolic-adaptation genes, including *ANLN*, *LOXL2*, *IGFBP2*, *ITGA5*, and *SLC2A3*. The favorable tail contained tumor-suppressive or lineage-restrictive genes, including *CBX7* (a component of Polycomb Repressive Complex 1), *TENT5C* (a non-canonical cytoplasmic poly-A polymerase frequently mutated in melanoma), and *CPEB3* (an RNA-binding protein that regulates translation and is a tumor suppressor specifically in hepatocellular carcinoma).

**Fig. 3:**
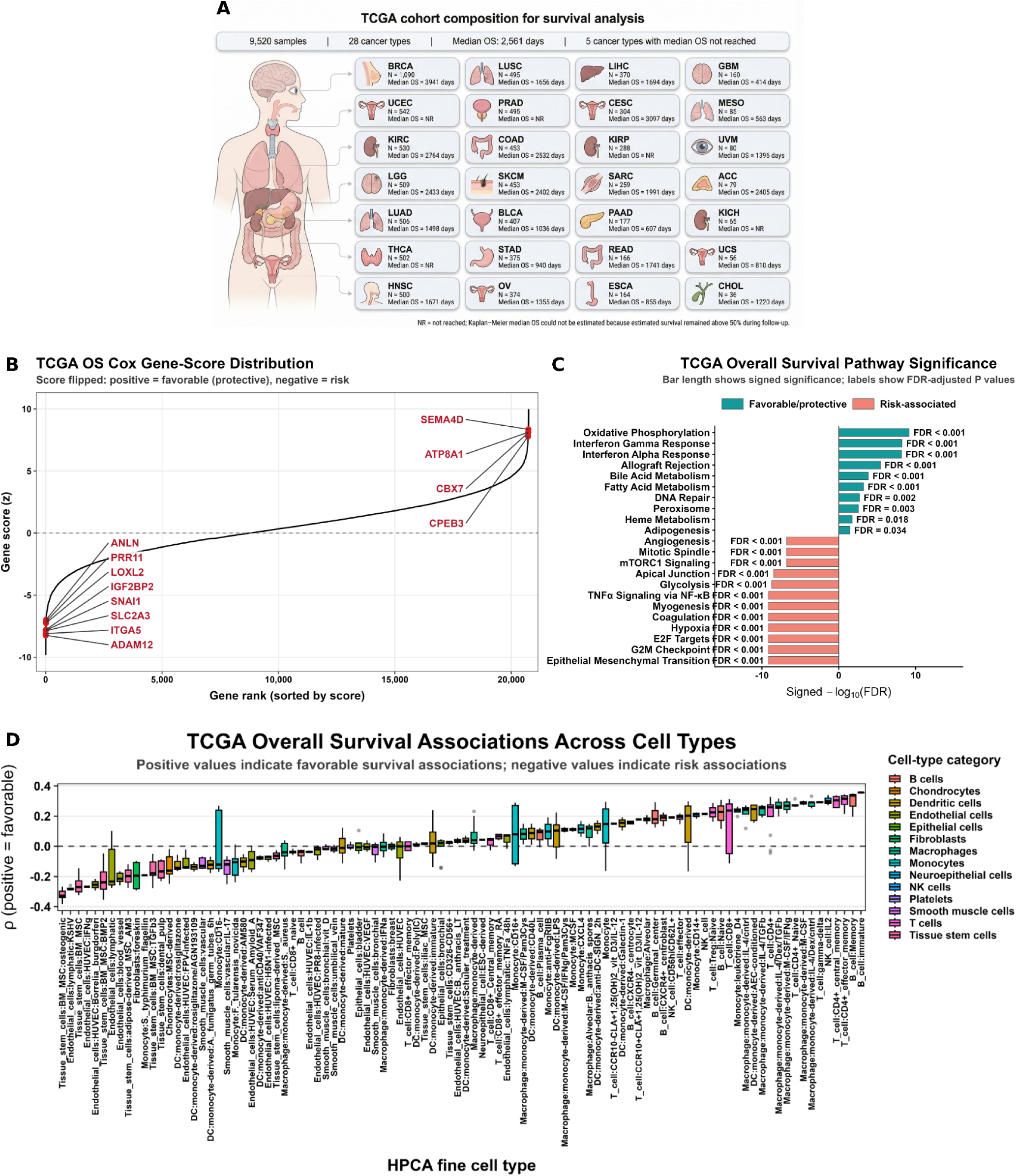
TCGA cohort composition and pan-cancer survival analysis identify favorable immune programs and mesenchymal risk states. (A) Composition of the TCGA cohorts used for pan-cancer overall survival (OS) analysis. (B) Ranked gene-level Cox OS scores identify genes associated with favorable versus unfavorable prognosis. (C) Hallmark GSEA shows immune-associated pathways on the favorable side and mesenchymal, hypoxic, angiogenic, and proliferative pathways on the risk side. (D) Projection of TCGA OS gene scores onto HPCA reference profiles shows that lymphoid programs, including γδ T-cells (6th from right), align with favorable prognosis.

Pathway analysis reinforced this favorable-versus-unfavorable structure. Better prognosis aligned with IFN-α response, allograft rejection, oxidative phosphorylation, DNA repair, and lipid-metabolic programs, whereas worse prognosis aligned with EMT, G2M checkpoint, E2F targets, angiogenesis, hypoxia, glycolysis, TNFα/NF-κB, mTORC1, coagulation, and myogenesis (Fig. 3C). Thus, favorable prognosis was associated not only with immune activation and inflammation but also with relative absence of hypoxic, glycolytic, angiogenic, and mesenchymal gene expression programs.

Projection of the TCGA overall survival gene scores onto the 157-population reference placed adaptive lymphoid lineages, including T, NK, and B cells, on the favorable side of the survival axis (Fig. 3D). Stromal, vascular, and tissue-stem-like signatures aligned with unfavorable prognosis. As in the ICB analysis, γδ T-cells were among the top favorable projected populations (ranked 6th), while monocyte and dendritic-cell states were more variable.

To test whether prognostic cell-state effects transfer across RNA detection technologies, we applied the same HPCA correlation-enrichment scorer to TCGA RNA-seq and 108 PRECOG (PREdiction of Clinical Outcomes from Genomic profiles) microarray cohorts comprising 13,116 patients (5). We fit a univariate Cox OS model per cohort × cell type and pooled within PRECOG cancer type by DerSimonian–Laird random-effects meta-analysis (Methods). For each of 17 PRECOG-vs-TCGA matched cancer pairs, we then computed Spearman correlation of the per-cell log-hazard ratios. The per-cell prognostic profiles were strongly concordant in cancers with dense coverage in PRECOG: lung adenocarcinoma (ρ = 0.97, 23 PRECOG cohorts), low-grade glioma (0.94, k = 12), melanoma (0.89, k = 3), liver (0.88), sarcoma (0.88) and bladder (0.83). The prognostic profiles remained positive in ovarian (0.73), prostate (0.72), head and neck (0.72), glioblastoma (0.56), gastric (0.30), pancreatic (0.29) and colon (0.21) cancer. Across the 17 matched pairs, 14 were sign-concordant (one-sided binomial p = 6.4 × 10⁻³).

### 3.4 ICB-associated cell-type programs show concordance with TCGA survival and recorded non-ICB treatment outcomes

We then asked whether the immune programs associated with ICB response were the same programs associated with favorable TCGA survival. The ICB-response and TCGA-OS cell-type rankings were positively correlated across HPCA cell types (R² = 0.495; p = 6.4 × 10⁻¹⁶; Fig. 4A), indicating that similar microenvironmental programs were associated with favorable outcomes in both settings.

**Fig. 4.**
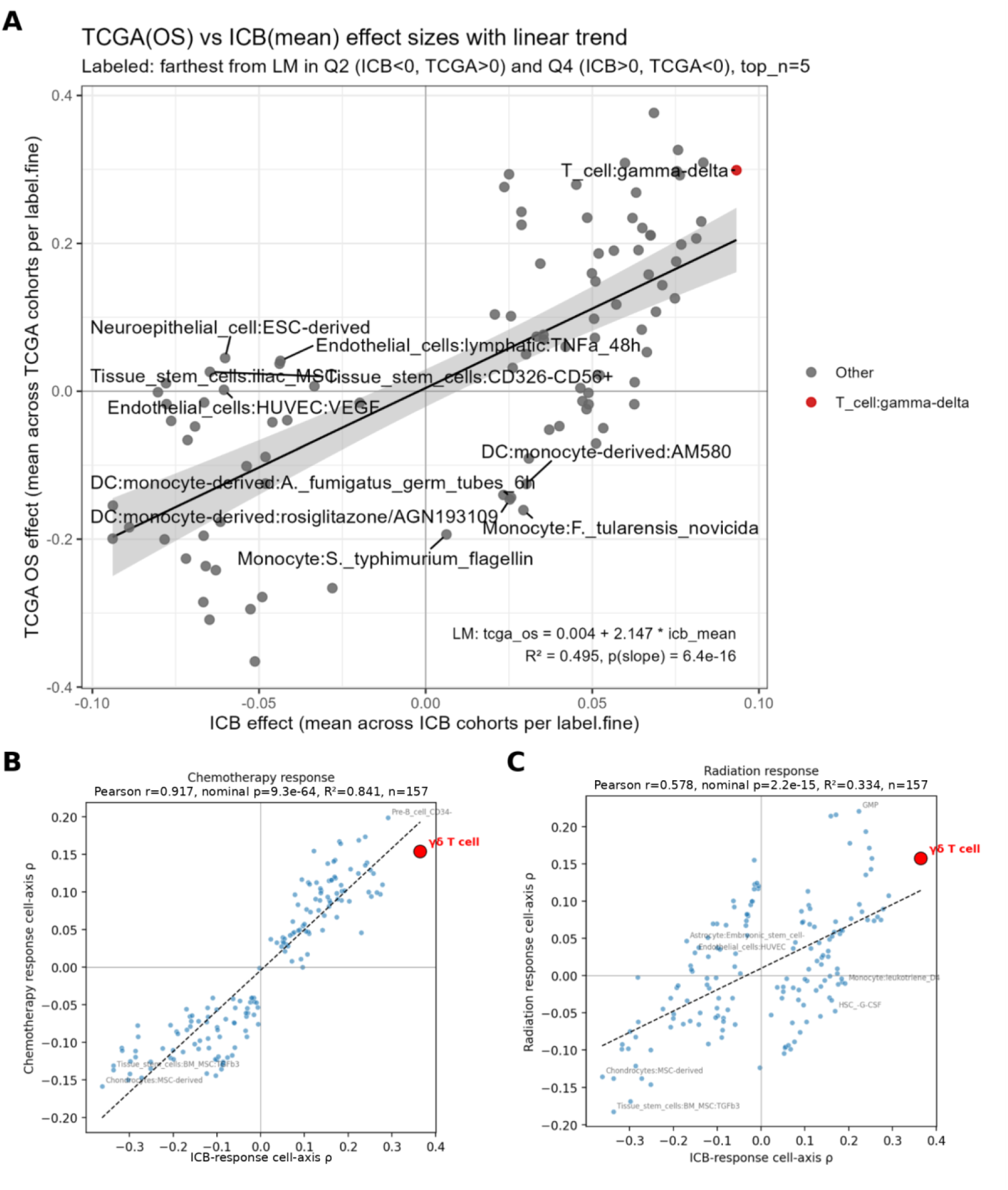
Cross-context concordance of ICB-favorable cell-type programs with TCGA survival and selected non-ICB treatment-response axes. **(A)** Cell-type effects from ICB response and TCGA overall survival are positively correlated, with γδ T-cells among the shared favorable populations. **(B)** The chemotherapy-response axis closely recapitulates the ICB-response cell-type landscape, with γδ T-cells remaining in the favorable quadrant. **(C)** The radiation-response axis shows moderate concordance with the ICB-response landscape, with γδ T-cells again aligned with the favorable quadrant.

The comparison also identified biologically interpretable discordance between ICB and other treatment types. Activated myeloid and antigen-presenting states were favorable for ICB response but unfavorable for TCGA overall survival, a pattern consistent with antigen-presenting-cell priming in ICB and chronic tumor-associated inflammation outside ICB (Fig. 4A). In contrast, vascular, stromal, and stem-like states were associated with reduced ICB responsiveness but had weaker associations with reduced TCGA OS, consistent with a stronger role for T-cell exclusion programs in checkpoint blockade response than in overall survival in TCGA setting, which includes mixed treatment exposures.

TCGA survival aggregates many treatment exposures. To test whether the favorable TME landscape applied to specific treatments, we built responder-vs-non-responder cell-type rankings for four non-immunotherapy TCGA treatment modalities with RECIST-evaluable records: chemotherapy (3,175 treatment lines from 1,406 patients across 25 cancer types), radiation (1,780 / 1,360 / 20), targeted therapy (153 / 124 / 8), and hormone therapy (115 / 85 / 2). Across the 157 HPCA cell types common to all axes, the ICB-response axis showed strong descriptive concordance with the chemotherapy-associated axis (Pearson r = 0.92, R² = 0.84, nominal p ≈ 1 × 10⁻⁶³; Fig. 4B) and moderate concordance with the radiation-associated axis (r = 0.58, R² = 0.33, nominal p ≈ 2 × 10⁻¹⁵; Fig. 4C). Concordance was weaker for targeted therapy (r = 0.26, R² = 0.07, nominal p = 1.1 × 10⁻³) and hormone therapy (r = 0.17, R² = 0.03, nominal p = 0.04), two strata with small sample sizes and narrower cancer coverage. Because the 157 HPCA profiles are correlated and share a common projection basis, these P values are nominal and descriptive.

γδ T-cells lay near the convergent core of these landscapes. On the ICB axis, the T_cell:gamma-delta reference ranked first among 157 cell types (ρ = +0.365). The same reference retained a favorable sign on the chemotherapy (ρ = +0.15) and radiation (ρ = +0.16) axes and showed inverse associations in the underpowered targeted (ρ = -0.14) and hormone (ρ = -0.07) strata. These data support γδ abundance as a marker of a favorable immune state in the three treatment settings with the greatest statistical power, while the targeted and hormone results should remain exploring.

### 3.5 Higher γδ T-cell abundance is associated with better ICB outcome and is not fully explained by CD8 abundance

Having found that γδ-associated programs mark the ICB responder axis, we asked whether per-patient γδ T-cell abundance in the TME tracked clinical outcomes. Across the patient-level ICB set (n = 1,356 with response; n = 1,074 with OS), pretreatment γδ TME abundance was associated with response and survival. The γδ score corresponded to an odds ratio of 1.38 for response (95% CI 1.23-1.56; p = 8.9 × 10⁻⁸; Fig. S3a) and a hazard ratio of 0.82 for OS (95% CI 0.76-0.88; p = 3.2 × 10⁻⁷; Fig. S3b). The direction of effect was consistent in 15 of 19 response cohorts (Fig. S5) and 11 of 13 OS cohorts (Fig. 5A). The opposite direction was observed in the ipi-progressed pre-treatment Riaz2017 cohort (17 patients after the split; p = 0.032). That reversal is not robust to the cut-point: the same cohort is not significant under a median split (p = 0.11) and in the continuous per-SD model (HR 1.14, 95% CI 0.76–1.70, p = 0.53).

**Fig. 5:**
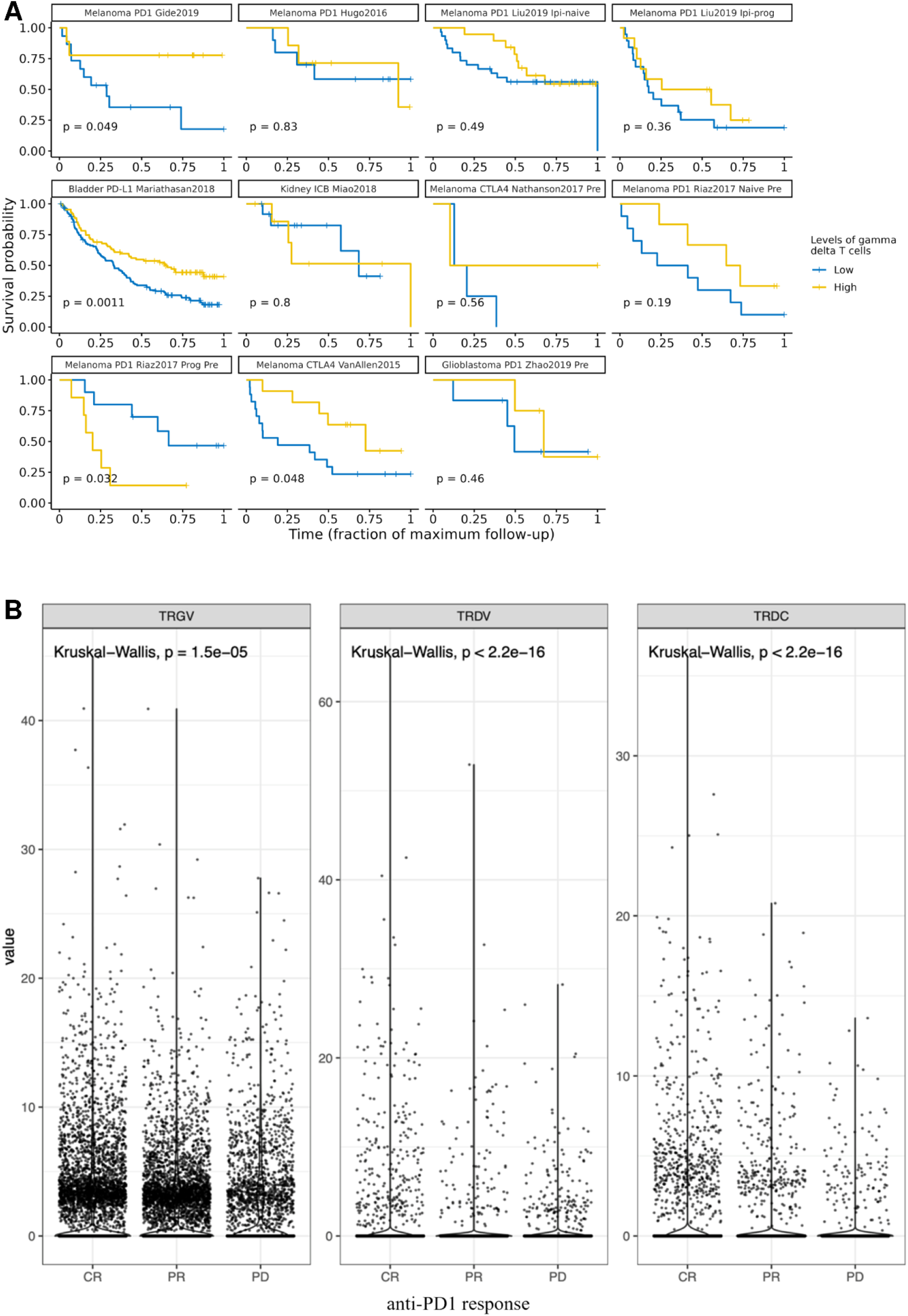
γδ T-cells are associated with improved outcomes in immunotherapy cohorts. (A) Kaplan-Meier survival curves across ICB cohorts stratified by high versus low γδ T-cell abundance, using a median split within cohort after standardization. (B) Single-cell analysis of γδ TCR gene expression across anti-PD-1 response categories (complete response (CR), partial response (PR), and progressive disease (PD)). Expression of γδ-associated TCR genes *TRGV*, *TRDV*, and *TRDC* differs across response groups, consistent with higher γδ TCR gene expression in better responders.

Because γδ abundance co-varies with broader T-cell inflammation, we next adjusted the ICB models for CD8 abundance using eight CD8 quantification strategies. Across these controls, the γδ response odds ratio remained between 1.28 and 1.54, with p < 1 × 10⁻⁴ in every model. The γδ OS hazard ratio remained between 0.80 and 0.86, with p < 1 × 10⁻³ in every model. These results indicate that CD8 adjustment did not remove the γδ association in the integrated ICB analysis (Fig. S3).

We then examined IMvigor210 (20), a urothelial cancer atezolizumab (anti-PD-L1 monoclonal antibody) cohort that classified tumors into three immune phenotypes (Inflamed, Excluded, Desert) based on CD8^+^ T-cell staining patterns visualized by immunohistochemistry (IHC), an approach orthogonal to RNA-seq. In this cohort (n = 347; IHC phenotype known for n = 283), the unadjusted γδ score was associated with response (OR 1.56; 95% CI 1.19-2.05; p = 1.5 × 10⁻³; Fig. S4a) and OS (HR 0.77; 95% CI 0.68-0.88; p = 1.1 × 10⁻⁴; Fig. S4b). Adjustment for the IHC immune phenotype gave similar estimates for γδ (response OR 1.56; 95% CI 1.15-2.13; p = 4.7 × 10⁻³; OS HR 0.78; 95% CI 0.67-0.90; p = 9.2 × 10⁻⁴; Fig S4a, b). Applying the same adjustment to the CD8 score itself left no significant CD8 effect (OR range 0.99-1.15; all p > 0.28; Fig S4c), supporting the phenotype as an effective CD8 control in this cohort.

For orthogonal support in immunotherapy-treated tumors, we examined γδ T-Cell receptor gene expression in a single-cell Hodgkin lymphoma dataset (Fig. 5B). Expression levels of the genes encoding T-cell receptor γ and δ chain variable and constant regions (*TRGV*, *TRDV*, and *TRDC*) were largely concordant with each other and showed concordant differences across complete response, partial response, and progressive disease categories, with higher expression in better-responding patients (Fig. 5B).

Together, these ICB analyses indicate that inferred γδ abundance carries response- and survival-associated information that is not removed by CD8 adjustment. The evidence remains associative, but it supports follow-up of γδ abundance as a candidate component of pretreatment immune profiling.

### 3.6 Higher γδ T-cell abundance is associated with better survival across several TCGA cancer types and is supported by receptor reconstruction

We next asked whether the association between γδ abundance in the TME and favorable outcome extended to tumors outside the ICB cohorts. Within each TCGA cancer type, we scored γδ T-cell abundance, split patients at the median, and compared OS. Higher γδ abundance was associated with better survival in several cancer types, most clearly melanoma, breast, ovarian, and bladder cancer (Fig. 6A). Prostate and kidney cancer showed little separation despite adequate sample size, and lung cancers showed only marginal separation.

**Fig. 6:**
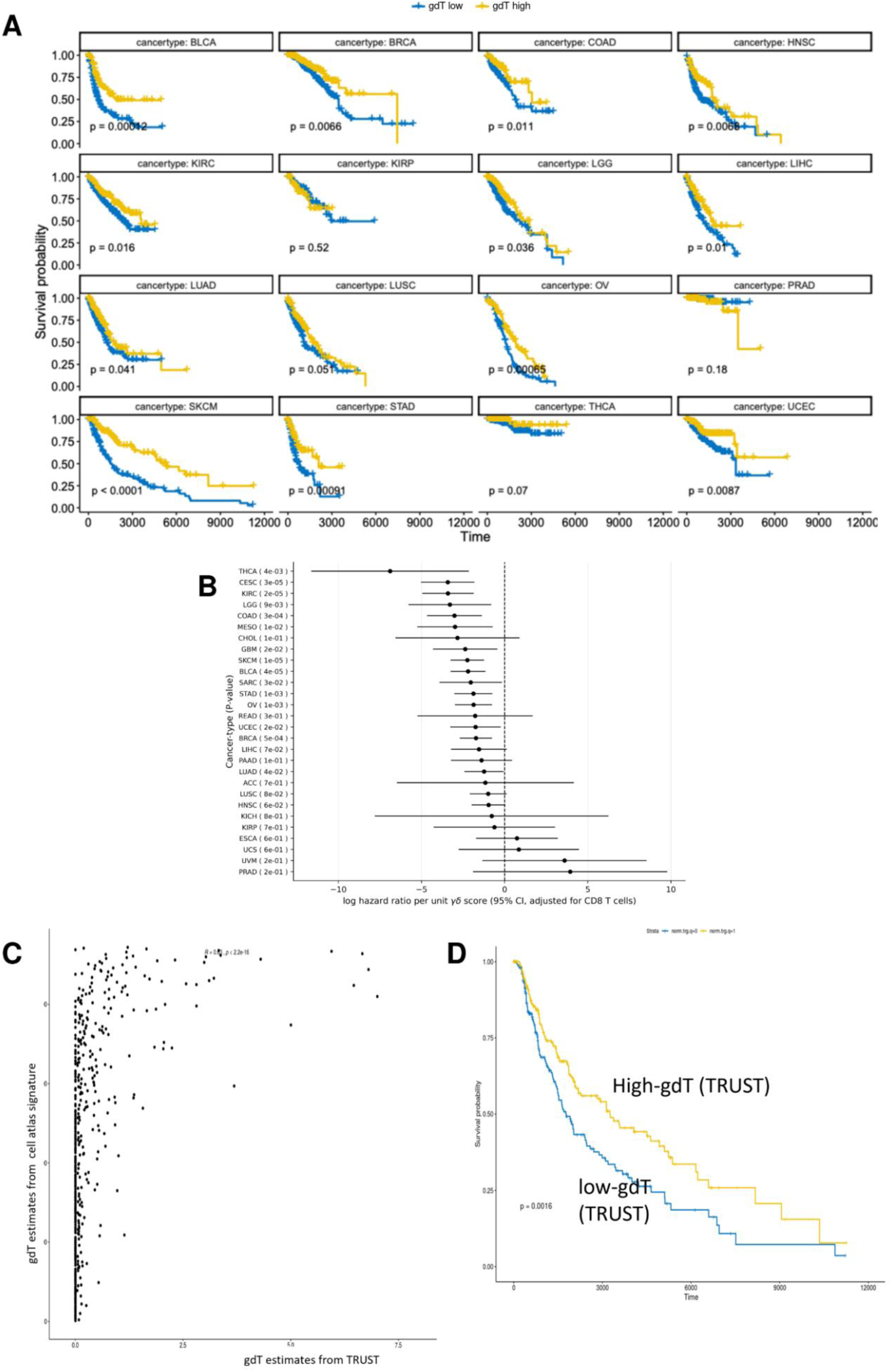
γδ T-cells show prognostic associations in TCGA and retain signal after CD8 adjustment. (A) Kaplan-Meier curves for OS across TCGA cancer types stratified by high (yellow) versus low (blue) γδ T-cell abundance. (B) Cancer-type-specific Cox log hazard ratio estimates for γδ abundance after adjustment for CD8^+^ T-cell abundance. P-values are indicated in the y-axis label. (C) Concordance between TRUST4-based γδ TCR reconstruction and the atlas signature-based γδ estimate. (D) Kaplan-Meier survival comparison between tumors with high versus low TRUST4-derived γδ TCR content.

These data are consistent with our previous CRISPR screen (36), which showed that unlike conventional CD8^+^ T-cells, γδ T-cells can recognize stress-induced ligands, phosphoantigens, butyrophilin molecules, and other nonclassical cues without strict MHC class I peptide presentation. Thus, γδ abundance is not simply a pan-immune-infiltration marker. The strongest associations were seen in melanoma, breast, ovarian, and bladder cancer, tumor types that frequently contain immune-inflamed and TLS/B-cell-rich subsets. By contrast, there was weaker separation in settings with low baseline immune activation (prostate cancer) or potentially dysfunctional immune infiltrates (e.g., renal cancer).

Because γδ abundance can co-vary with cytotoxic inflammation, we adjusted the TCGA survival models for CD8^+^ T-cell abundance. After CD8 adjustment, the γδ survival association remained evident in multiple cancer types (Fig. 6B), indicating that the TCGA signal is not wholly attributable to CD8-associated inflammation.

We then asked whether the γδ abundance signal in the TCGA data depended on the method used to infer γδ abundance by transcriptional similarity to a reference atlas. Instead, we used TRUST4-based de novo reconstruction of TRG/TRD receptors to produce receptor-based estimates of γδ abundance. These new estimates correlated with the signature-based scores (R = 0.43; p < 2.2 × 10⁻¹⁶; Fig. 6C). Tumors with higher reconstructed γδ TCR content also showed better survival (log-rank p = 0.0016; Fig. 6D). These data support γδ abundance as a reproducible prognostic correlate in TCGA rather than a feature of any single quantification method.

Within the cross-platform replication of cell-type rankings established in section 3.3, γδ T-cells were showed favorable associations on both platforms in 14 of 17 matched cancers (one-sided binomial p = 6.4 × 10⁻³) and reached significance on both platforms in head and neck (HR = 0.59, p = 0.014), melanoma (0.82, p = 0.04), ovarian (0.86, p = 6 × 10⁻⁶), and colon cancer (0.78, p = 8 × 10⁻⁴).

## 4. Discussion

Across pretreatment ICB cohorts and TCGA, γδ T-cell abundance and γδ-associated transcriptional programs were reproducible correlates of favorable outcomes, with effect sizes that varied by tumor type, cohort, and treatment context. ICB responders tended to have pre-treatment tumor microenvironments with interferon-rich, cytotoxic, and lymphoid-enriched expression programs that included γδ T-cell signatures. In TCGA samples across cancer types and treatment modalities, related lymphoid programs aligned with favorable survival. The ICB-favorable cell-type landscape was strongly concordant with chemotherapy response and moderately shared with radiation response. These findings extend earlier studies that linked leukocyte subsets to prognosis and ICB benefit (5, 6) and are also concordant with reports linking γδ-associated transcriptional programs to clinical benefit (9–11). Our analysis differs from these earlier studies in scope. Previous pan-cancer immune-deconvolution studies catalogued broad leukocyte–prognosis associations (5, 6) and recent γδ-focused work established γδ effector activity in defined single contexts (HLA-class-I–deficient tumors (37) and a pan-cancer γδ TCR repertoire (38)). In contrast, we evaluate γδ abundance jointly across immunotherapy and non-immunotherapy modalities and test whether its outcome association persists after CD8 adjustment. Our analysis identifies γδ T-cells as a recurrent correlate of a favorable pretreatment immune state in the TME but does not establish it as a standalone causal determinant.

The MHC-independent role of γδ T-cells in tumor control offers a mechanistic framework for the association with positive outcomes. γδ T-cells can recognize malignant or stressed epithelium through multiple receptor-ligand axes, including NKG2D ligands and butyrophilins (39), providing routes for MHC-independent tumor recognition. In this way, γδ T-cells may help control tumors that have escaped classical CD8^+^ T-cell surveillance through loss or downregulation of MHC class I, for example, via β2-microglobulin/MHC I pathway defects (2).

We also identified several candidate modifiers of the relationship between γδ T-cell abundance and outcome. First, we observed cooperation between γδ T-cells and B-cell/TLS biology, with some of the strongest survival associations in tumors that are both γδ T-high and Memory-B/TLS-high. This is consistent with a model in which γδ T-cells promote or benefit from organized intratumoral humoral immunity. Second, high fibroblast/stromal signatures attenuate the favorable association with γδ T-cell abundance, suggesting physical or cytokine-mediated barriers (for example, TGF-β-rich stroma) that warrant further mechanistic study. Third, epithelial differentiation programs, for example, keratinization and ELF5-high states, appear to support γδ T benefit, implying tumor-cell-intrinsic ligands or stress programs that modulate γδ T recognition. These phenotype-by-context effects are correlative and identify candidate modifiers including B-cell/TLS organization, stromal density, and epithelial state. Any causal contribution of these modifiers requires prospective testing. Consistent with this context dependence, prior disease-specific studies have reported favorable γδ T-cell associations in dMMR tumors (9), cervical cancer (14), Merkel cell carcinoma (10), and breast cancer (11, 12), contrasted with less favorable or even adverse γδ associations in some head and neck squamous cell carcinoma cohorts (16).

Several considerations support, but do not prove, that the γδ signal is partially independent of the CD8-dominated T-cell program. Adjustment for CD8 abundance (gene-expression-based, deconvolution-based, and IHC-based) attenuates but does not abolish the γδ association in ICB cohorts and TCGA (forest plot, Fig.6B and Fig. S3). These data support prospective evaluation of whether pretreatment γδ T-cell abundance provides incremental information beyond CD8-centric ICB biomarkers. γδ T-cells are currently being advanced into the clinic as adoptive cellular products and engineered CAR therapies (13, 39, 40). Prospective trials should determine whether γδ T-cell abundance is useful for stratification or enrichment in studies of γδ T-cell–based cellular therapies or γδ T-cell agonists, alone or in combination with immune checkpoint blockade.

Several limitations constrain interpretation. Bulk RNA-seq cannot resolve γδ T-cell subtypes such as Vδ1/Vδ2 or cytotoxic versus IL-17-skewed states (which may be pro-tumorigenic). The TCGA modality analyses linked primary-tumor RNA-seq with recorded treatment outcomes that could occur later in the clinical course, and some patients contributed multiple treatment records. Consequently, these analyses do not establish a treatment-specific predictive effect and should be interpreted only as exploratory concordance between tumor-expression programs and recorded outcomes. TRUST4 reconstruction of γδ T-cell receptors provided support in TCGA, but repertoire reconstruction has not yet been extended across all ICB cohorts. Clinical endpoints, treatment exposures, and sample sizes vary across both the ICB and TCGA cohorts, which limits direct cross-cohort comparison and motivates random-effects pooling rather than fixed estimates. Pipeline calibration was not deliberately conservative and was performed under a single simulation design. Under the global null, the joint multivariable model yielded a γδ type-I error of 0.04 at α = 0.05 and the approximately 0.10 rate was observed only in the marginal one-cell-at-a-time model. Therefore, to assess the robustness of this finding, we relied on cross-platform PRECOG replication and orthogonal TRUST4 reconstruction rather than nominal p-values alone. Concordance between our RNA-seq-based data and microarray-based PRECOG data was most pronounced in lung and melanoma and weaker in other cancers. However, weaker agreement in those cancers reflects smaller PRECOG sample numbers and probe/platform differences and therefore should not be read as the absence of a γδ effect in those tumors. TCGA treatment classifications rely on recorded treatment fields and keyword matching, which might be noisy or incomplete. Targeted and hormone response axes are underpowered.

This study provides evidence that γδ T-cell abundance in the pretreatment TME is an indicator of favorable cancer-treatment outcomes in both immunotherapy and non-immunotherapy settings. The γδ signal is complementary to CD8 metrics and appears to be modulated by B-cell/TLS organization, stromal architecture, and the degree of epithelial-to-mesenchymal transition (EMT). This work suggests several future studies, including prospective validation in uniformly annotated pretreatment cohorts, extension of TRG/TRD reconstruction to ICB datasets, and single-cell/repertoire studies that resolve γδ subtypes and their interactions with B cells, fibroblasts, and tumor epithelium.

## Ethics Statement

All data analyzed in this study were obtained from publicly available, de-identified sources. No new patients were enrolled, and no new human or animal experimentation was conducted; therefore, no additional institutional ethics approval or informed consent was required for this study.

## Supplementary figures

**Fig. S1.**
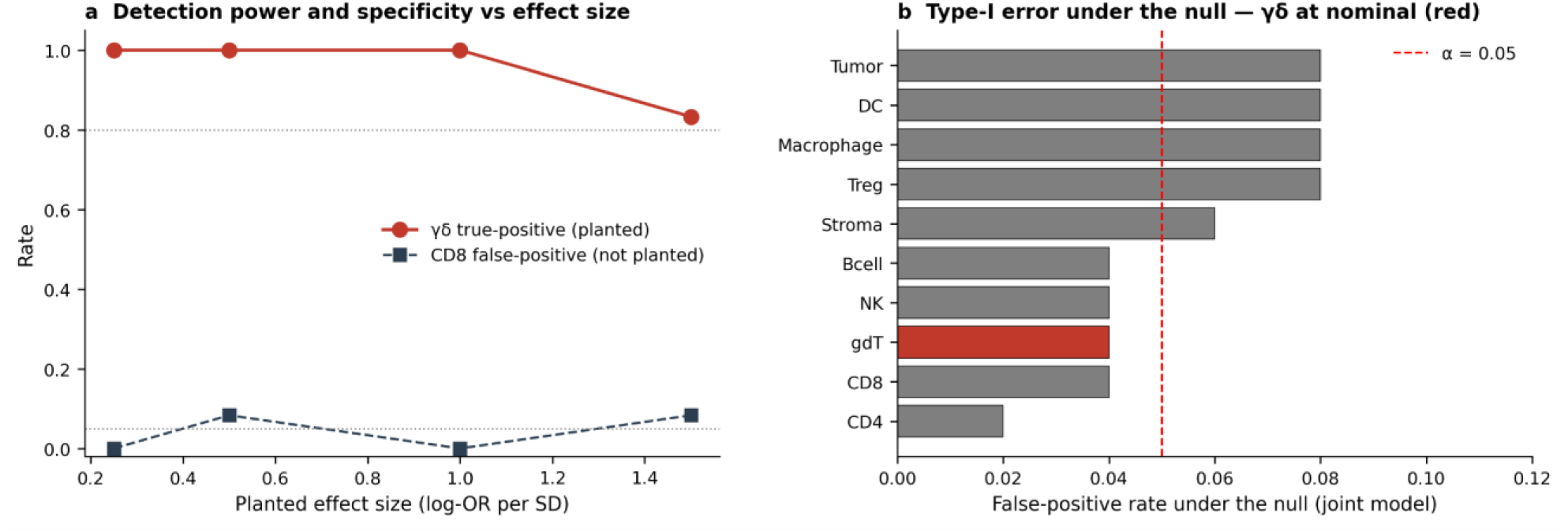
Pipeline calibration (joint multivariable model). (a) Detection power and specificity versus planted effect size: simulated γδ effects are recovered at approximately 100% power across realistic effect sizes, while the false-positive rate for a non-planted CD8 signature stays near α. (b) Type-I error under the global null: the γδ false-positive rate is at the nominal level (0.04 at α = 0.05), with no γδ-specific inflation. Pseudo-bulk simulation design in Methods §2.4.

**Fig. S2.**
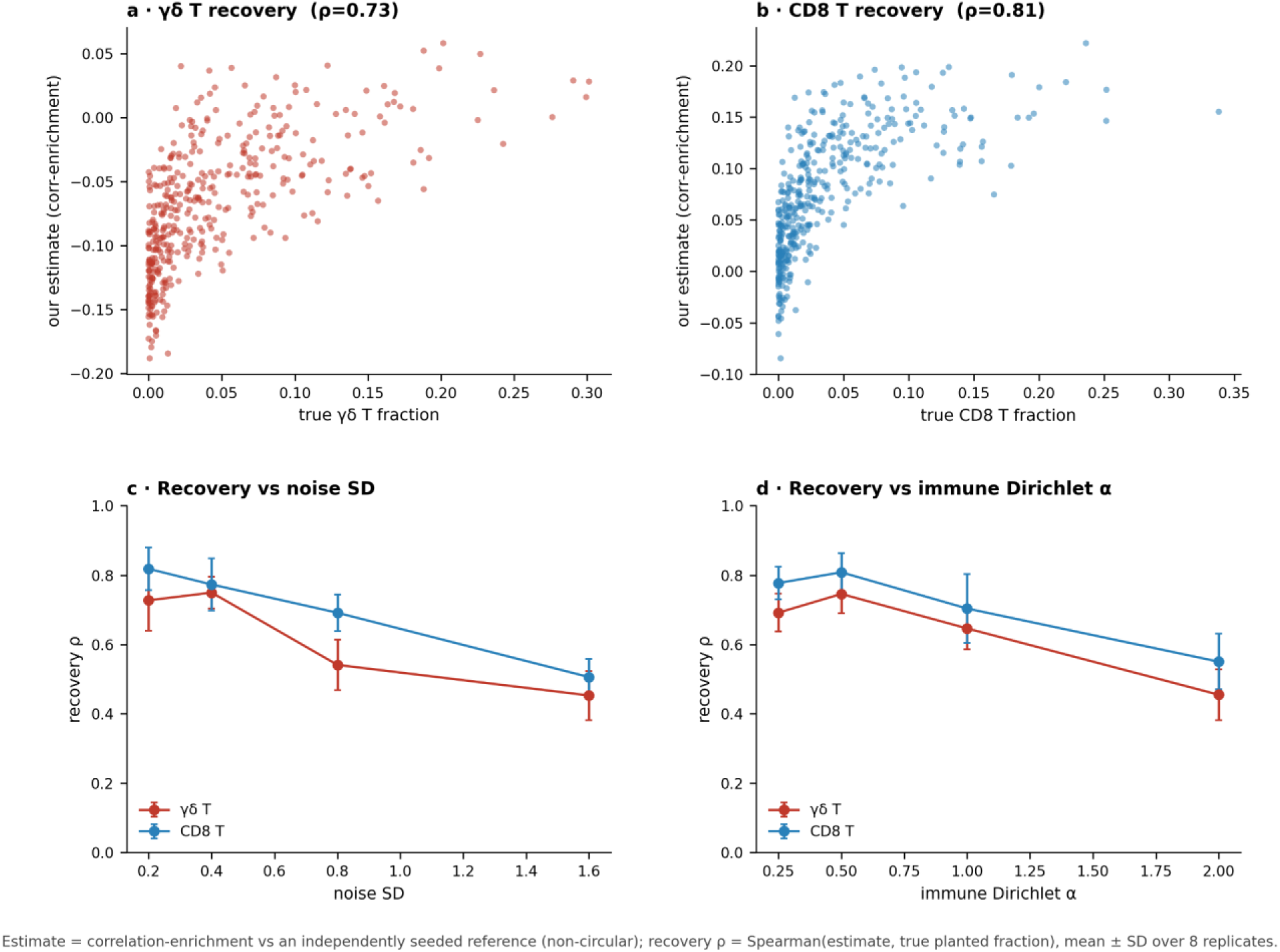
Per-sample abundance scores recover ground-truth cell fractions in simulation. Pseudo-bulk samples were built from known cell-type mixtures and scored by correlation-enrichment against an independently seeded reference (non-circular). (a, b) Recovery of the planted γδ and CD8 T-cell fractions (Spearman ρ = 0.73 and 0.81). (c, d) Recovery as a function of added noise (c) and mixture concentration (Dirichlet α; d); values are mean ± SD over 8 replicates. Methods §2.4.

**Fig. S3.**
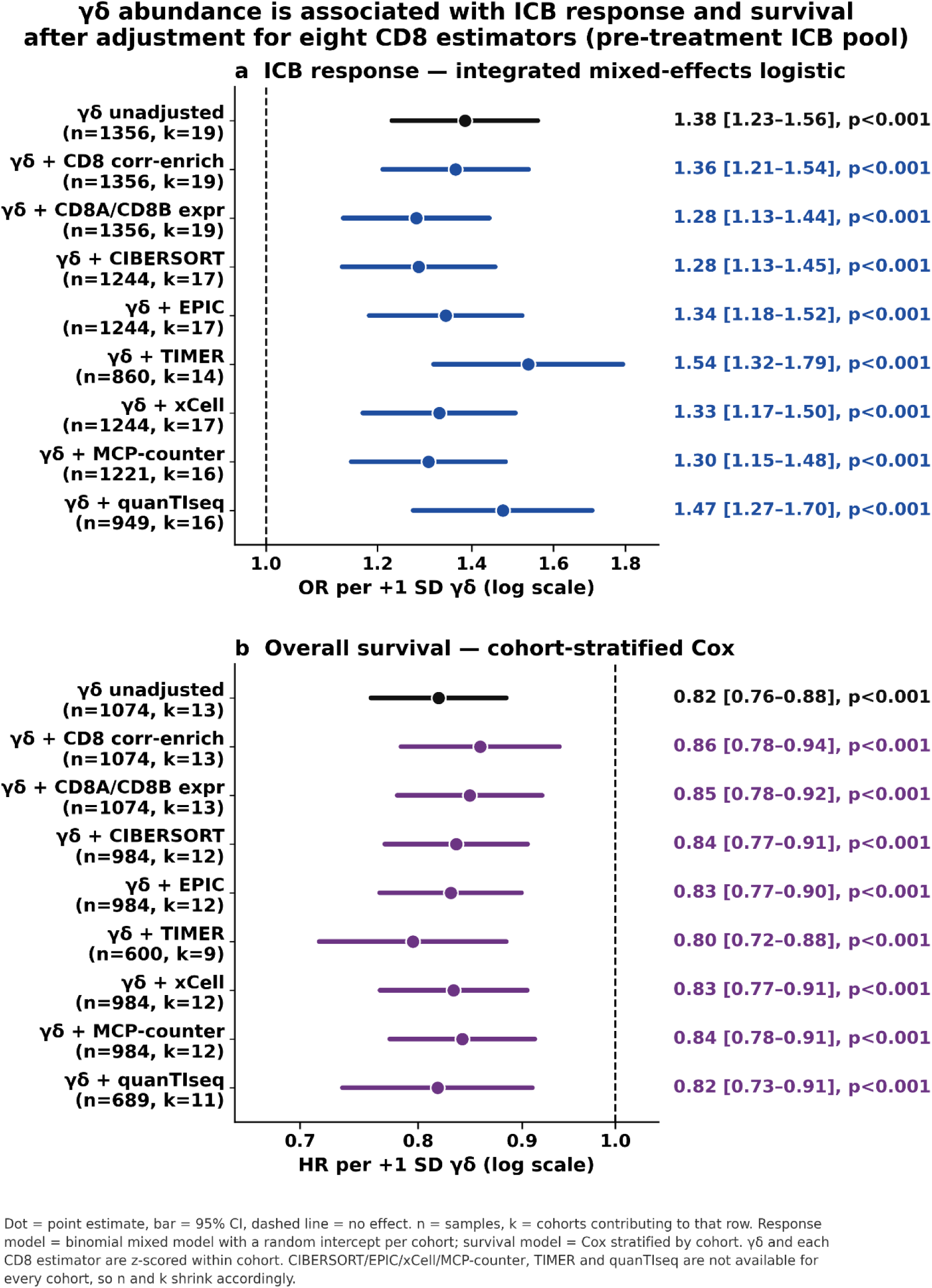
γδ abundance is associated with ICB response and survival after adjustment for eight CD8 estimators. Pooled analysis of pretreatment ICB cohorts (19 cohorts, n = 1,356 for response; 13 cohorts, n = 1,074 for OS). (a) ICB response (binomial mixed-effects logistic): γδ odds ratio per +1 SD, shown unadjusted (top row) and after adding each of the eight CD8 estimators as a covariate; all remain significant (OR 1.28–1.54, p < 0.001). (b) Overall survival (cohort-stratified Cox): γδ hazard ratio per +1 SD, unadjusted and CD8-adjusted (HR 0.80–0.86, p < 0.001). γδ and each CD8 estimator are z-scored within cohort; CIBERSORT/EPIC/xCell/MCP-counter, TIMER, and quanTIseq are not available for every cohort, so n and k vary by row. Methods §2.4.

**Fig. S4.**
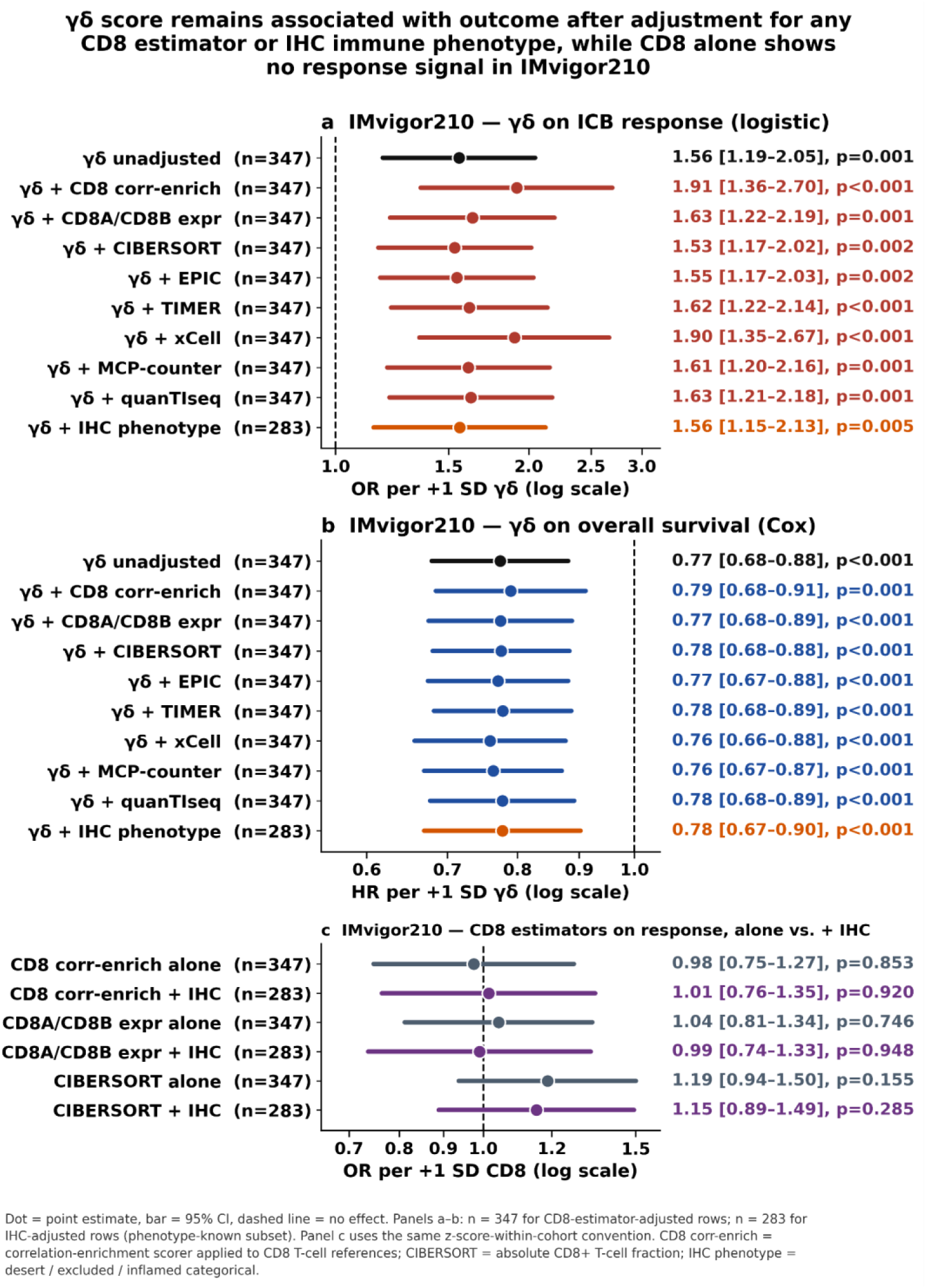
In IMvigor210, the γδ–outcome association remains after adjustment for each CD8 estimator and the orthogonal IHC immune phenotype, while CD8 alone is uninformative. Atezolizumab-treated urothelial cohort (n = 347; IHC immune phenotype available for n = 283). (a) ICB response (logistic): γδ odds ratio per +1 SD, unadjusted (top row) and after adding each of eight CD8 estimators or the IHC phenotype as a covariate; all remain above 1 and significant (OR 1.53–1.91; every p < 0.005). (b) Overall survival (Cox): γδ hazard ratio per +1 SD, unadjusted and after the same adjustments (HR 0.76–0.79; every p < 0.005). (c) Specificity control: each CD8 estimator was fitted as the predictor in place of γδ, alone and after IHC-phenotype adjustment; CD8 carries no response signal (OR 0.98–1.19; all p > 0.15), so the γδ effect is not a relabeled CD8 effect. γδ and CD8 are z-scored within cohort. Methods §2.4.

**Fig. S5.**
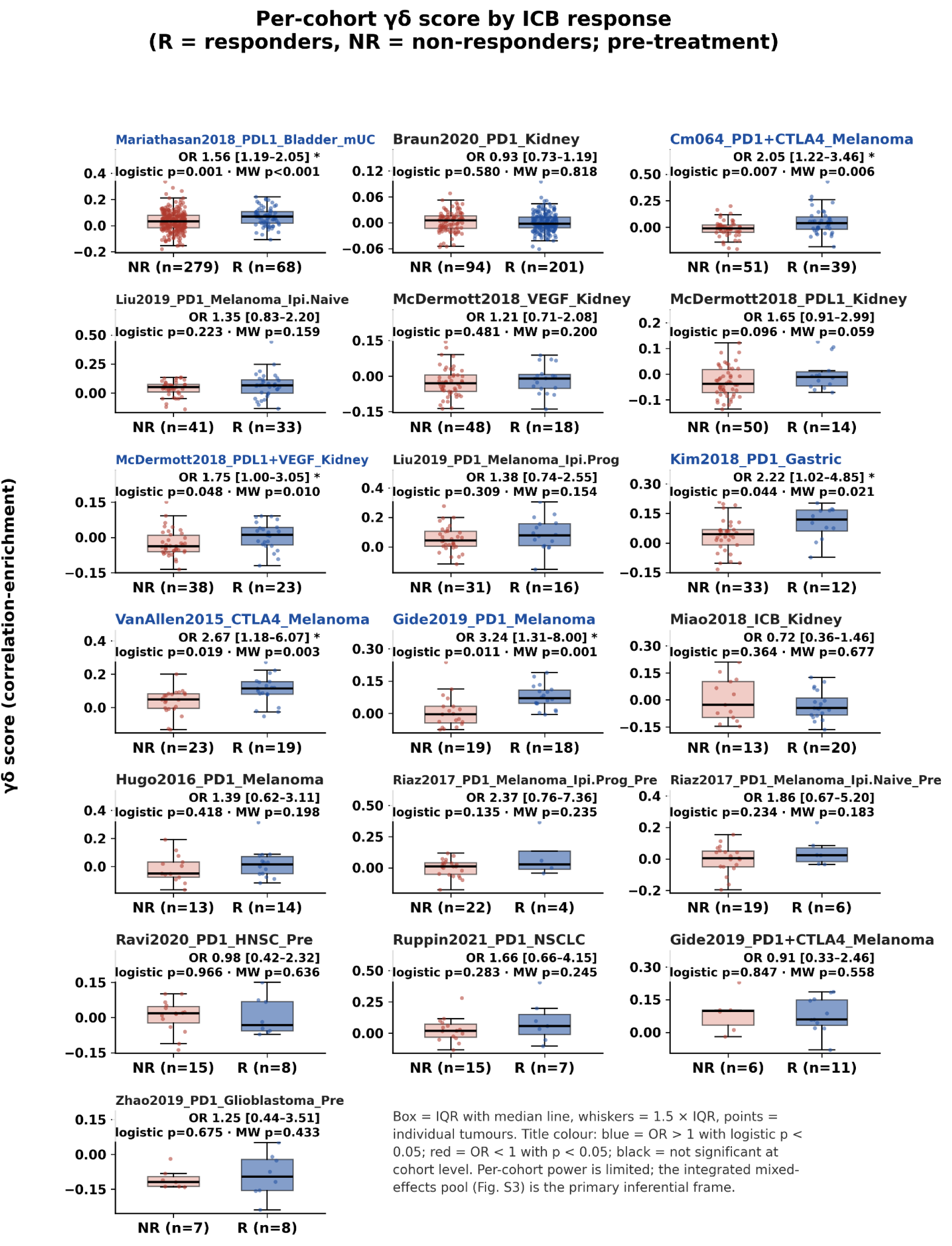
Per-cohort γδ score by ICB response. Per-cohort distribution of the per-sample γδ correlation-enrichment score in non-responders (NR) versus responders (R), one panel per ICB cohort with response data (19 cohorts, pretreatment). Each title gives that cohort’s logistic odds ratio (per +1 SD γδ) and Mann–Whitney p; blue titles mark a significant favorable association (OR > 1, p < 0.05), black titles denote nonsignificant results. γδ was higher in responders (OR > 1) in 15 of 19 cohorts, reached per-cohort significance in 6 (Mariathasan, CheckMate-064, Gide PD-1, Van Allen, Kim, McDermott PD-L1+VEGF), and no cohort showed a significant adverse association; individual cohorts are underpowered, so the integrated mixed-effects pool (Fig. S3) is the primary inferential framework. Methods §2.4.

